# A soybean pattern recognition receptor conferring broad-spectrum pathogen and pest resistance regulates expression of several NLR receptor proteins

**DOI:** 10.1101/2023.04.04.535565

**Authors:** Micheline N. Ngaki, Subodh K. Srivastava, Wang Feifei, Madan K. Bhattacharyya

**Affiliations:** Department of Agronomy, Iowa State University, Ames IA 50011; Department of Entomology and Plant Pathology, North Carolina State University, Raleigh, NC, 27695, USA; Northeast Institute of Geography and Agroecology, Key Laboratory of Soybean Molecular Design Breeding, the Chinese Academy of Sciences, Harbin 150081, China

**Keywords:** soybean, *GmDR1*, chitin, RNA-seq, plant basal resistance, NB-LRR, transcriptomics, RT-qPCR

## Abstract

Overexpressors of *Glycine max disease resistant 1* (*GmDR1*) exhibits broad-spectrum resistance against *Fusarium virguliforme*, soybean cyst nematode (SCN), spider mites, and soybean aphids in soybean. To understand the mechanisms of broad-spectrum immunity mediated by *GmDR1*, we investigated the transcriptomes of a strong and a weak *GmDR1*-overexpressors following treatment with chitin, a pathogen- and pest-associated molecular pattern (PAMP) found in these organisms. The strong and weak *GmDR1*-overexpressors exhibited altered expression of 6,098 and 992 genes, respectively, as compared to the nontransgenic control following chitin treatment. However, only 192 of these genes exhibited over two-fold changes in expression levels in both strong and weak *GmDR1-*overexpressors as compared to the control. MapMan analysis of the 192 genes revealed 64 biotic stress-related genes, of which 53 were induced and 11 repressed as compared to the control. The 53 chitin-induced genes include nine that encode receptor kinases, 13 encode <u>n</u>ucleotide-binding leucine-rich <u>r</u>epeat (NLR) receptor proteins, seven encoding WRKY transcription factors, four ethylene response factors, and three MYB-like transcription factors. Investigation of a subset of these genes revealed three receptor protein kinases, seven NLR proteins, and one WRKY transcription factor genes that are induced following *F. virguliforme* and SCN infection. The integral plasma membrane GmDR1 protein most likely recognizes PAMPs including chitin and activates transcription of genes encoding receptor kinases and NLR proteins. GmDR1 could be a pattern recognition receptor that regulates the expression of several NLRs for expression of PAMP-triggered immunity and/or priming the effector triggered immunity.

## Introduction

In the early step of plant defenses, molecular patterns associated with the invading pathogen and pests are recognized by the plant plasma membrane-localized pattern-recognition receptors (PRRs) including receptor-like kinases (RLKs) or receptor-like proteins (RLPs) [1]. It has been documented that PRRs recognize microbe- (or pathogen-) associated molecular patterns (MAMPs or PAMPs) including the bacterial flagellin, elongation factor-Tu and peptidoglycan, and the fungal polysaccharide chitin and the oomycete glucan to activate the first layer of plant immunity known as pattern-triggered immunity (PTI) [2, 3].

Among the plant PAMPs, the fungal cell wall feature and linear polymer of N-acetyl-D-glucosamine chitin is an important activator of the plant immunity and defense mechanisms [4-6]. PAMPs such as chitin and glucan are liberated by the plant chitinases and glucanases through degradation of fungal walls during pathogen invasion [6, 7]. Multiple reactions are induced by these cell wall components including the expression of defense-related genes following recognition of these PAMPs by the PAMP recognition receptors [7, 8].

The second layer of defense, the effector-triggered immunity (ETI), is regulated by intracellular <u>n</u>ucleotide-binding domain and leucine-rich <u>r</u>epeat (NLR) containing receptors termed resistance (R) proteins that recognize the pathogen effector proteins [1, 9, 10]. Plant genome contains hundreds of genes encoding NLR proteins that play an important role in the perception of pathogen effectors and conferring race- or gene-specific disease resistance [11-13]. The nucleotide-binding (NB) domains bind nucleotides and contain highly conserved motifs including kinase-2, P-loop, GLPL or Gly-Leu-Pro-Leu, MHD motifs [14-16]. The leucine-rich repeat (LRR) domain is involved in effector recognition that determine the expression of race- or gene-specific resistance [17]. The N-terminal region of NLR proteins possess a Toll/interleukin-1 receptor (TIR) domain in TIR-NLR (TNLR) proteins or a coiled-coil (CC) domain in CC-NLR (CNLR) proteins [18, 19].

Despite innumerable studies have dissected PTI, ETI, and their downstream signaling pathways regulated by the plant hormones, salicylic acid (SA), jasmonate (JA), and ethylene (ET), much is unknown about the mechanisms of broad-spectrum resistance against multiple pathogens and pests regulated by PAMP recognition receptors. In the United States, over a dozen pathogens routinely cause annual soybean yield suppression valued close to $4 billion [20]. The soybean cyst nematode (SCN) annually suppresses soybean yield valued at over $1.1 billion and *Fusarium virguliforme* over $0.4 billion [20]. *F. virguliforme* causes sudden death syndrome (SDS) and rapidly suppresses accumulation of transcripts of a limited number genes including *Glycine max disease resistance 1* (*GmDR1*) [21]. Its overexpression enhances resistance of soybean against the SCN (*Heterodera glycines*), and two insect pests, spider mites (*Tetranychus urticae*, Koch) and soybean aphids (*Aphis glycines*, Matsumura) in addition to *F. virguliforme* [22]. This study suggested that *GmDR1*-mediated broad-spectrum resistance could be initiated through perception of the PAMP chitin found in all four soybean pathogen and pests. Chitin has been demonstrated to interact with receptor complex CERK1 and LYK5 RLKs [23, 24]. In rice, the RLP chitin-elicitor binding protein CEBiP recognizes chitin [5]. In soybean, the transmembrane cysteine-rich receptor-like kinases (CRKs) involved in plant immunity binds to chitin [25]. In this study, we applied a transcriptomic approach to investigate the molecular basis of broad-spectrum resistance mediated by overexpression of *GmDR1*. We identified 64 biotic stress-related related genes, of which 53 are induced and 11 repressed by chitin. The chitin-induced genes include 13 NLR-type genes including 10 TNLR, three CNLR, seven WRKY transcription factors, four ethylene response factors, and three myeloblastosis (MYB)-like proteins. We have shown that a subset of these genes is induced following infection of soybean with *F. virguliforme* and SCN suggesting that *GmDR1* is a master-regulator of several NLR-type disease resistance genes, which could govern the expression of broad-spectrum disease resistance in soybean.

## Results

### Investigation of the transcriptomes regulated by *GmDR1*

To investigate the molecular basis of broad-spectrum disease and pest resistance mediated by *GmDR1* against two pathogens, *F. virguliforme* and SCN, and two pests, spider mites and soybean aphids, we compared the transcriptomes of two *GmDR1* overexpressors: (i) weak overexpresser, the DR1-136 line and (ii) strong overexpresser, the DR1-107 line with that of the (iii) nontransgenic control Williams 82 line following treatment with the PAMP chitin [22]. We investigated the transcriptomes of the *GmDR1*-overexpressors following chitin treatment to determine the role of *GmDR1* on the global defense gene regulation.

The overview of the steps taken in generating RNA-seq data is summarized in Fig. S1A. Young leaves were sampled at two different time-points, 12 and 24 h following treatment with either chitin or phosphate buffer used to dissolve chitin as a negative control from three independent experiments. To determine levels of gene expression, the reads were mapped to individual genes of the soybean genome sequence (*Glycine max* Wm82.a4.v1), with the cutoff value was set to fragments per kilobase of transcript per million mapped reads (FPKM). Before initiating analyses of the raw data, we investigated if the treated soybean stem cut seedlings responded to chitin appropriately. We were able to identify previously reported chitin-responsive genes [26] that were induced 12 h following treatment of nontransgenic Williams 82 line with chitin (Table S1).

The gene expression patterns following chitin and buffer treatments were complex. More genes were transcriptionally regulated in the strong *GmDR1*-overexpresser DR1-107 as compared to the weak *GmDR1*-overexpresser DR1-136 following chitin treatment when compared to the nontransgenic Williams 82 control (Fig. S1B; Dataset S1). Although both transgenic lines confer resistance against the fungal pathogen *F. virguliforme* [22], the number of differentially expressed genes (DEGs) in the strong *GmDR1*-overexpresser (DR-107) was much higher as compared to that in the weak *GmDR1*-overexpresser (DR1-136) line when the gene expression levels of the two lines were compared with that of the nontransgenic Williams 82 line 12 h following chitin or buffer treatment (Fig. 1A; Dataset S1). By 24 h, the number of DEGs between weak *GmDR1*-overexpresser line DR1-136 and nontransgenic Williams 82 control line was only three as opposed to 7,596 between the strong *GmDR1*-expresser line DR1-107 and Williams 82 (Fig. 1B).

**Figure 1.**
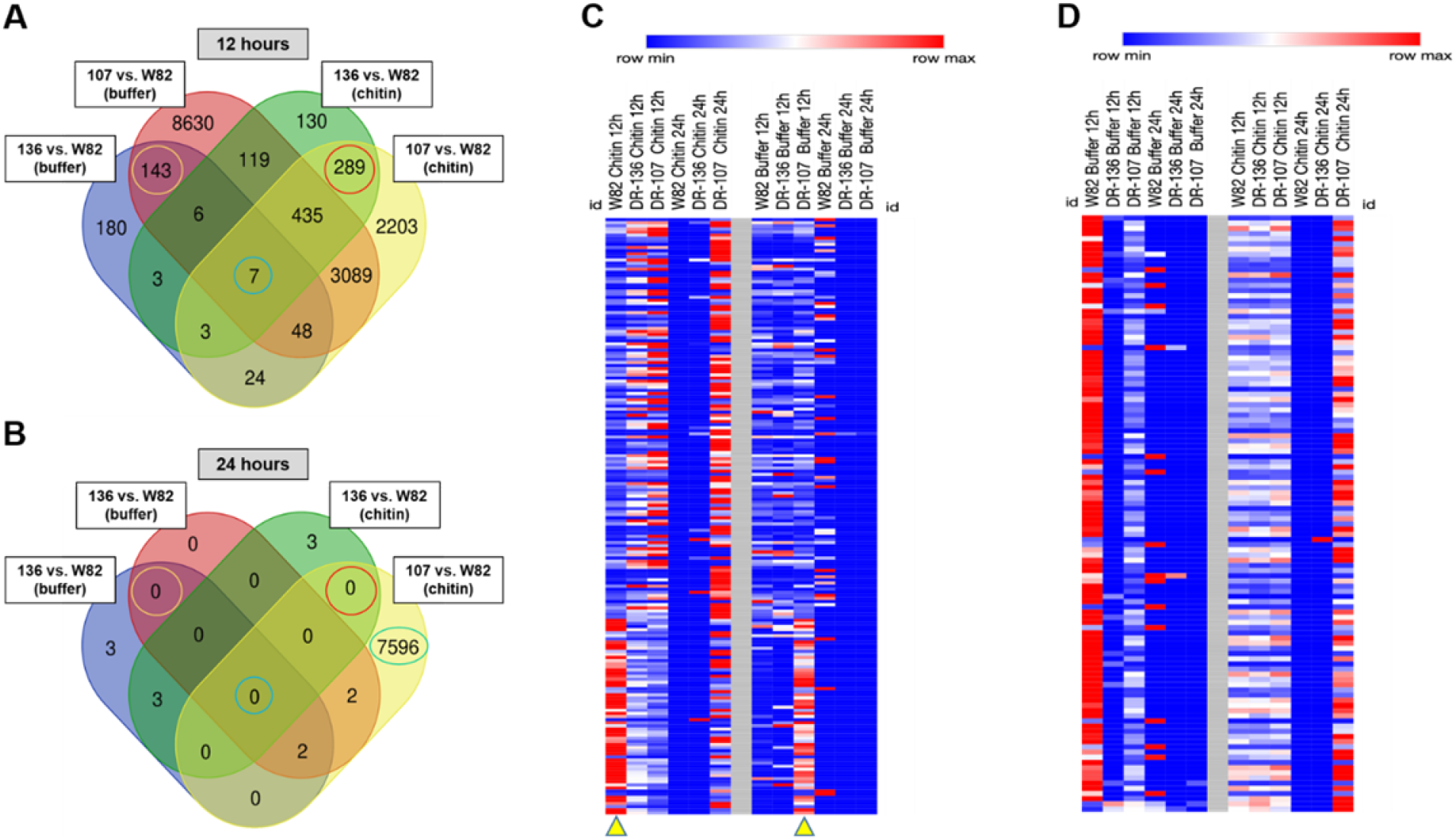
Frequency and expression patterns of differentially expressed genes (DEGs). *(A)* equencies of genes that are differentially expressed in the weak (136, DR1-136) and strong 07, DR1-107) *GmDR1*-overexpresser as compared to the nontransgenic Williams 82 (W82) ne 12 h following chitin or phosphate buffer (buffer) treatment. Note that 143 (shown with ange circle) and 289 (red circle) DEGs are common to both *GmDR1*-overexpressers 12 h llowing buffer and chitin treatments with at least 1.5-fold change and *p* ::;*0*.*05* between illiams 82 and each of the *GmDR1*-overexpressers 12 h following chitin treatment Expression seven genes (shown with green circle; Table S2) was significantly altered in *GmDR1*-erexpressers 12 h following chitin and buffer treatments. *(B)* Frequencies of genes that are fferentially expressed in the weak (136, DR1-136) and strong (107, DR1-107) *GmDR1*-erexpressors as compared to the nontransgenic W82 line 24 h following chitin or phosphate ffer treatment. Note that no DEGs were detected in both *GmDR1*-overexpressors 24 h llowing chitin or buffer treatment. *(C)* Heat map displaying the differential expression patterns the 192 (129 Induced and 69 repressed, Dataset S2) of the 289 chitin-responsive DEGs that ere differentially expressed at least by 2-fold and *p* ::;0.001 between each of the *GmDR1*-erexpressors and W82 12 following chitin treatment. Differential expression levels of these nes 24 h following chitin treatment, and 12 and 24 h following buffer treatment are also esented. *(D)* Heat map displaying the differential expression patterns of the 115 (2 Induced and 3 repressed, Dataset S3) of the 143 buffer-responsive DEGs that were differentially expressed least by 2-fold and *p* ::;0.001 between each of the *GmDR1*-overexpressors and W82 12 llowing buffer treatment. Differential expression levels of these genes 24 h following buffer treatment, and 12 and 24 h following chitin treatment are also presented.

### Identification of several *GmDR1*-regulated genes associated with the expression of broad-spectrum pathogen and pest resistance

Inclusion of the strong and weak *GmDR1*-overexpressors conferring broad-spectrum pathogen and pest resistance allowed us to identify a subset of only 289 genes that were differentially expressed in both *GmDR1*-overexpressors as compared to the nontransgenic control line 12 h following chitin treatment (Fig. 1A). Of these 289 genes, only 192 showed at least ≥ 2-fold changes in transcript levels (*p* value ≤ 0.001) in both overexpressors as compared to that in nontransgenic Williams 82 control (Dataset S2).

We also observed an influence of the overexpressed *GmDR1* gene on the differential expression of 143 genes 12 h following phosphate buffer treatment (Fig. 1A). However, only 115 genes of the 143 genes showed at least ≥ 2-fold changes in transcript levels (*p* value ≤ 0.001) in both overexpressors as compared to the nontransgenic Williams 82 control following phosphate buffer treatment (Dataset S3).

By 24 h following chitin and phosphate buffer treatments, no DEG was observed between nontransgenic control and both overexpressors (Fig. 1B). We observed only three DEGs in weak *GmDR1*-expresser and over 7,000 DEGs in the strong *GmDR1*-expresser, not common to both overexpressors, 24 h following chitin treatment, when compared transcript levels with that of the nontransgenic control (Fig. 1B). We identified seven genes that are differentially regulated between nontransgenic control and either of the overexpressors 12 h but not 24 h following chitin and phosphate buffer treatments (Fig. 1A; Table S2).

Heatmaps of the selected 192 and 115 genes differentially expressed between the *GmDR1* overexpressors and nontrangenic control 12 h following treatment with chitin and phosphate buffer, respectively, revealed that of the 192 chitin-responsive DEGs, 129 (67%) were induced, whereas 63 (33%) were repressed in both transgenic lines as compared to the nontransgenic control (Fig. 1C-D, Dataset S2). Interestingly, 113 of the 115 buffer-responsive DEGs were repressed and only two were induced among the *GmDR1*-overexpressors as compared to Williams 82 following buffer treatment (Dataset S3).

### Identification of key genes transcriptionally regulated by overexpressed *GmDR1*

To identify the defense-related genes and their relative transcript abundance regulated by *GmDR1*, the MapMan program was applied to investigate the selected subsets of 192 and 115 genes showing > 2-fold changes in transcript levels in both *GmDR1*-overexpressors as compared that in the nontransgenic Williams 82 control line 12 h following chitin and buffer treatment, respectively. Investigation of the 192 chitin-responsive DEGs revealed 64 biotic stress-related and 18 metabolism pathways-related genes (Fig. 2A; Dataset 4A-B). Of the 64 biotic stress-related genes, 53 were upregulated and 11 were downregulated following chitin treatment. Of the 53 upregulated genes, 23 genes encode 10 receptor kinases and 13 NLR proteins (11 TNLR and two CNLR), seven genes encode WRKY transcription factors implicated in resistance of soybean against its pathogens [27, 28]; four encode ethylene response factors (ERFs) modulating biotic and abiotic stresses [29], and three encode MYB-like proteins [30] (Table 1; Dataset 4A). Among the 115 buffer-responsive DEGs, two abiotic stress-related, 26 biotic stress-related, and seven metabolism pathway-related genes were identified (Fig. 2B; Dataset 4C-D).

**figure 2.**
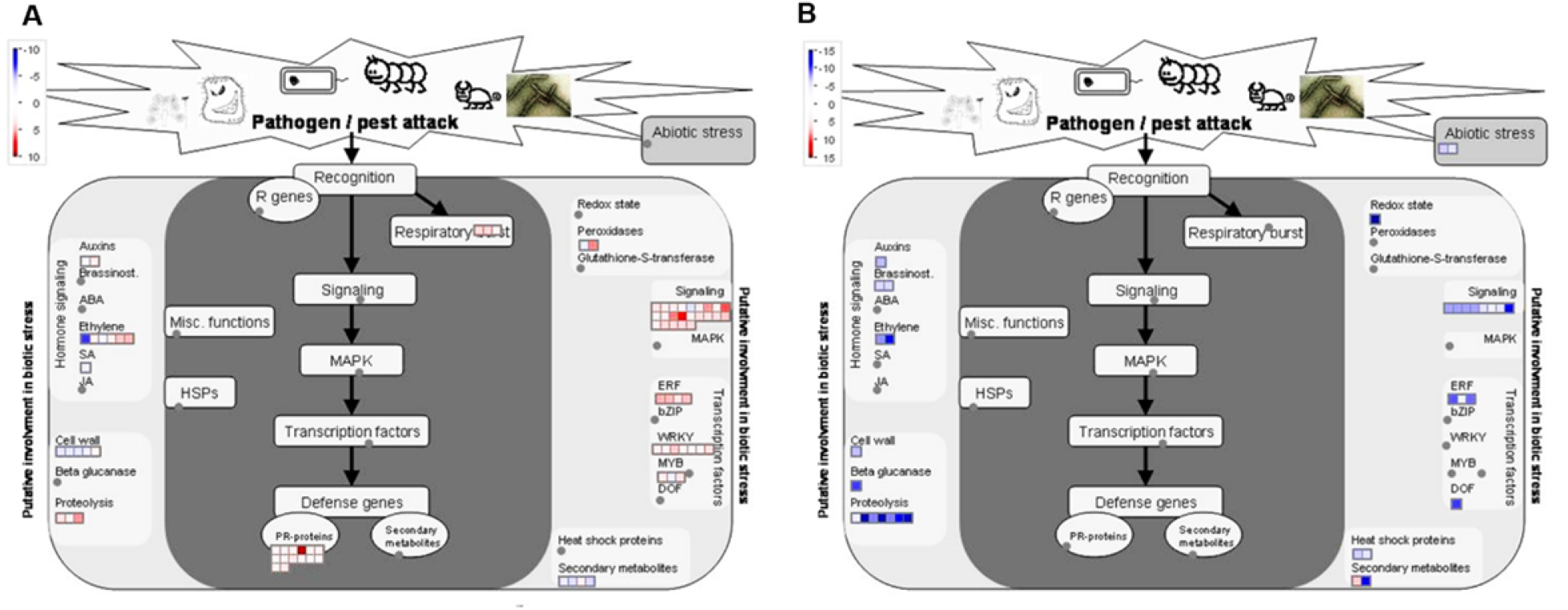
Genes regulated by overexpressed *GmDR1* following treatment with chitin (A) or hosphate buffer (B). The 192 chitin-responsive and 115 buffer-responsive DEGs common to th strong and weak *GmDR1* overexpressors (Fig.1) were visualized using the MapMan software. *(A)* From the 192 chitin-responsive genes, 64 involved in the biotic-stress pathways ataset S4A) and 18 in metabolic pathways (Dataset S4B) were identified. *(B)* From the 115 ffer-responsive genes, 28 involved in the abiotic and biotic-stress pathways (Dataset S4C) and seven in metabolic pathways (Dataset S4D) were identified. Blue indicates a decrease, whereas red indicates an increase (log2 fold changes) in the transcription levels of a gene when compared ith that of the nontransgenic Williams 82 control.

**Table 1.** Expression levels of the 22 DEGs encoding nine LRR receptor-like kinases and 13 NLR-type disease resistance proteins 12 h following chitin or buffer treatment.

| Gene name | Annotation | C3 | C4 | C5 | C6 | C7 |
| --- | --- | --- | --- | --- | --- | --- |
| *LRR-receptor kinases* |  |  |  |  |  |  |
| Glyma.05G126400 | LRR Transmembrane Protein Kinase | 3.1 | 3.3 | -2.1 | 1.1 | -1.1 |
| Glyma.14G118800 | LRR receptor-like Ser/Thr- Protein Kinase | 4.3 | 11.8 | -9.2 | -1.3 | -2.0 |
| Glyma.16G182700 | LRR Receptor-Like Protein Kinase | 5.5 | 5.3 | 1.0 | 1.0 | 1.2 |
| Glyma.16G182866 | LRR receptor-like Ser/Thr- Protein kinase GSO1 | 4.7 | 4.1 | 1.0 | 1.0 | 1.4 |
| Glyma.16G184200 | LRR Receptor-Like Protein Kinase | 2.1 | 2.7 | -1.3 | 1.3 | 1.7 |
| Glyma.16G186100 | LRR Receptor-Like Protein Kinase | 3.0 | 9.9 | 1.2 | -1.1 | -1.2 |
| Glyma.16G186600 | LRR Receptor-Like Protein Kinase | 3.3 | 2.7 | -1.1 | 1.2 | 1.3 |
| Glyma.16G193600 | LRR Receptor-Like Protein Kinase | 2.3 | 3.5 | -1.9 | 1.6 | 1.8 |
| Glyma.18G254000 | LRR Receptor-Like Protein Kinase | 3.3 | 2.9 | -1.4 | -1.2 | -1.0 |
| *NBS-LRR-type Disease Resistance Receptors* |  |  |  |  |  |  |
| Glyma.07G077700 | NB-ARC-LRR, RPS2-like | 2.2 | 2.2 | -1.3 | -1.1 | -1.2 |
| Glyma.17G180300 | CC-NBS-LRR, RPW domain | 2.4 | 2.8 | 1.8 | 1.8 | 1.3 |
| Glyma.16G137000 | TIR-NBS-LRR class | 2.9 | 4.0 | 1.0 | 1.0 | -1.1 |
| Glyma.16G137600 | TIR-NBS-LRR class | 2.6 | 3.3 | -2.1 | 2.0 | 1.9 |
| Glyma.13G028100 | TIR-NBS-LRR class | 2.4 | 3.1 | 1.1 | 1.5 | 1.5 |
| Glyma.16G136600 | TIR-NBS-LRR class | 2.3 | 4.1 | 1.5 | 1.4 | 1.2 |
| Glyma.11G153000 | TIR-NBS-LRR class | 2.1 | 2.7 | -5.9 | -1.0 | 1.1 |
| Glyma.12G135600 | TIR-NBS-LRR class | 2.0 | 2.1 | -2.2 | 1.4 | 1.5 |
| Glyma.03G048500 | TIR-NBS-LRR, RPS4-like | 5.0 | 3.1 | -1.0 | 2.1 | 2.0 |
| Glyma.03G048200 | TIR-NBS-LRR | 5.6 | 4.8 | 1.0 | 1.2 | 1.2 |
| Glyma.09G075500 | TIR-NBS-LRR, RPS4-like | 2.1 | 2.1 | -2.9 | 1.2 | 1.0 |
| Glyma.03G053500 | TIR-NBS-LRR | 30.1 | 9.5 | 1.0 | 1.1 | 1.1 |
| Glyma.03G053602 | TIR-NBS-LRR | 18.8 | 8.0 | 1.0 | 1.3 | 1.2 |
| *Signaling Proteins* |  |  |  |  |  |  |
| Glyma.17G011700 | F-box family protein with a domain of unknown function | 2.6 | 2.3 | 2.2 | 2.2 | 1.2 |
| Glyma.11G003500 | Pyridoxal-5'-phosphate-dependent enzyme family protein | 5.3 | 43.3 | -2.1 | -1.3 | -2.8 |
| Glyma.10G148700 | Calmodulin-binding protein | 2.9 | 5.8 | -2.4 | -2.4 | 1.4 |
| Glyma.04G223200 | WRKY DNA-binding protein 55 | 2.6 | 6.1 | -4.5 | -1.3 | -1.6 |
| Glyma.04G057700 | Integrase-type DNA-binding superfamily protein | 3.1 | 6.2 | 1.6 | 1.0 | -1.3 |
| Glyma.03G066800 | NAD(P)-linked oxidoreductase superfamily protein | 2.2 | 2.3 | 2.6 | 2.0 | 2.1 |

C3, Column 3: Fold changes (FCs) in DR1-136 Vs W82 12 h following chitin treatment
C4, Column 4: FCs in DR1-107 Vs W82 12 following chitin treatment
C5, Column 5: FCs in DR1-136 Vs W82 12 h following buffer treatment
C6, Column 6: FCs in DR1-107 Vs W82 12 h following buffer treatment
C7, Column 7: FCs in W82 12 h following chitin Vs buffer treatment

Among the chitin-induced DEGs, we identified seven defense-related WRKY transcription factor genes, the transcription levels of which were altered (Dataset S2 and S4). In plants, defense mechanisms are modulated by the expression of multiple transcription factors including WRKY proteins, ET-responsive factors (ERFs), and MYB-like proteins [28, 30, and 31]. It was reported that soybean genes encoding WRKY transcription factors were differentially expressed following *Phytophtora sojae* infection [32, 33]. Similarly, *F. graminearum* infection induced the expression of more than 10 *BdWRKY* genes in *Brochypodium distachyon* [34]. *WRKY22* is upregulated by chitin in Arabidopsis [35]. Expression of five *GmWRKY* genes enhanced SCN resistance in soybean roots [27]. Interestingly, among the seven DEGs encoding GmWRKY chitin-induced transcription factors identified in this study, *Glyma*.*03G042700* and *Glyma*.*04G223200* were shown to be upregulated following *P. sojae* and *Peronospora manshurica* infection, respectively [33, 36]. We discovered through RT-qPCR that *Glyma*.*04G223200* is induced in roots of *GmDR1*-overexpressors following *F. virguliforme* and SCN infection (Fig. S2, Dataset S2). Our study also identified four chitin-induced DEGs encoding ERF proteins and three encoding MYB-like proteins.

Here again, our data is supported by previous studies that support plant ERF transcription factors are involved in biotic and abiotic defense responses [37, 38]. In soybean, *GmERF113* is important for resistance against *P. sojae* [29]; and it was shown that the overexpression of *GmERF3* enhances abiotic stress tolerance and disease resistance in tobacco [39]. It was also shown that the overexpression of *GmERF5* induces immunity of soybean against *P. sojae* [40]. Furthermore, MYB-like proteins are regulators of biotic and abiotic defense responses [30, 41]. Thus, characterization of more transcription factors will likely contribute to generate disease resistant crop plants.

The following five highly chitin-induced genes among *GmDR1-*overexpressors belong to protein families putatively involved in plant immunity (Table S3). (1) *Glyma*.*11G003500* (FC = 25) encoding a pyridoxal-5-phosphate-dependent enzyme family protein with sequence-specific DNA binding domains is putatively involved in biosynthesis of tryptophan, the precursor of biologically essential metabolites that are involved in plant defenses [42]. 2) *Glyma*.*03G068200* (average FC = 21) encoding a drought-repressed protein is involved in soybean-aphid interactions [43]. (3) *Glyma*.*03G053500* (average FC = 20) encoding an uncharacterized TNLR protein. (4) *Glyma*.*03G053602* (FC = 13) encodes an uncharacterized TNLR protein. (5) *Glyma*.*05G141200* (FC = 13) encoding a jasmonate-zim-domain (JAZ)-like protein involved in the modulation of plant-pathogen interactions [44]. On the contrary, the five most highly chitin-repressed DEGs encode two proteins of unknown functions, a feruloyl CoA ortho-hydroxylase, a citrate-binding protein, and a mitochondrial-processing peptidase (Table S4).

This study also revealed that feeding of phosphate buffer through cut stems of soybean seedlings regulates many genes. Surprisingly, of the 115 DEGs 113 were suppressed and two were induced in the *GmDR1*-overexpressors as compared to Williams 82 12 h following buffer treatment (Table S5, Dataset S3). This observation suggest that GmDR1 also recognizes damage-associated molecular patterns (DAMP) released from the cut soybean stem and/or buffer-associated cues.

### Identification of the subset of chitin-responsive genes differentially regulated following *F. virguliforme* and *H. glycines* infection

The differential regulation of 23 genes encoding LRR domain containing receptor-like proteins and other immunity-related signaling genes led us to wonder if any of these genes have putative roles in conferring *GmDR1*-mediated broad-spectrum resistance against pathogens and pests. As a first towards answering the relevance of these chitin-induced receptors and signaling genes in broad-spectrum resistance, we conducted quantitative reverse transcriptase polymerase chain reaction (RT-qPCR) to determine if the expression levels of 28 genes including 13 NLR, nine receptor-kinases, and six candidate genes for signaling and metabolic pathways are altered in soybean roots following infection with either *F. virguliforme* or SCN. The expression data revealed that six TNLR, one CNLR, three receptor protein kinases, and one WRKY transcription factor gene were induced in roots of at least the strong *GmDR1*-overexpresser, DR1-107 following *F. virguiforme* and SCN infection (Fig. 3A-B; Table S6).

**Figure 3.**
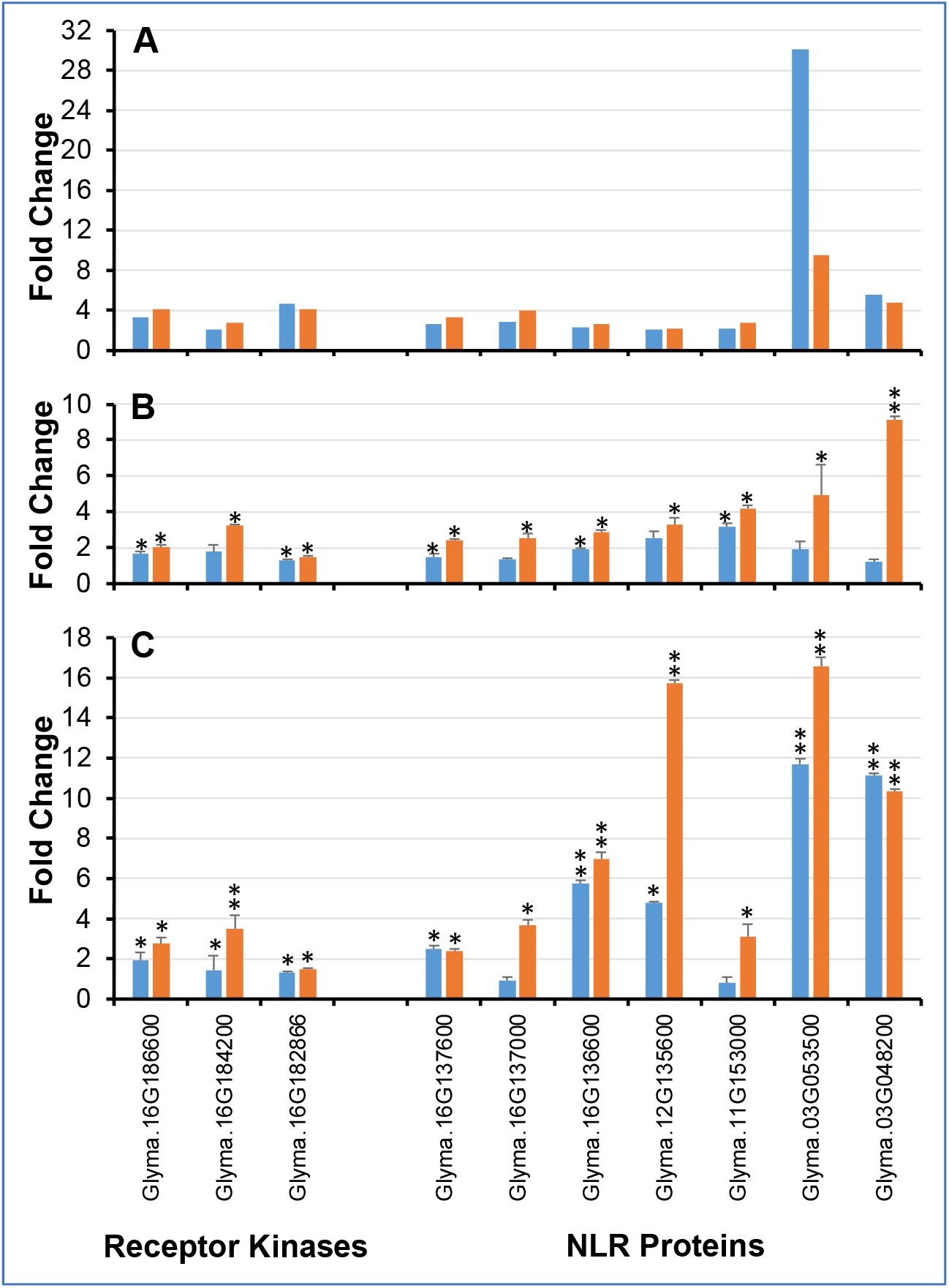
A subset of chitin-responsive genes encoding NLR proteins and LRR-receptor kinases e induced in *GmDR1*-overexpressors following *F. virguiforme* and SCN infection. *(A)* gnificantly enhanced expression levels of LRR-receptor kinases and NLR genes 12 h following itin treatment (RNA-seq data). Fold changes were calculated by comparing FPKM values of a ne in *GmDR1*-overexpressors, either DR-107 or DR1-136, with that in the nontransgenic ntrol Williams 82 line 12 h following chitin treatment. Transcription fold changes of the genes ere at least 2-fold and *p* ≤.0.05. *(B)* Significantly enhanced expression levels of receptor nases and NLR genes 12 days following *F. virguliforme* infection. Fold changes of the genes tween *GmDR1*-overexpressors and Williams 82 following *F. virguliforme* infection were termined by RT-qPCR. Data in *B* and *C* are mean ± SE determined from three biological plications. The soybean *Elongation factor 1-b* (*ELF1-b*) gene (*Glyma*.*02g44460*) was used as internal control in RT-qPCR. *, *p* ≤ 05 and **, *p* ≤ 01. *(C)* Significantly enhanced expression vels of receptor kinases and NLR genes 21 days following SCN infection. Fold changes of the nes between *GmDR1*-overexpressors and Williams 82 following SCN infection were termined by RT-qPCR. Data for all 28 DEGs selected for RT-qPCR are presented on Fig. S2.

### Gene ontology (GO) functional classification of DEGs

To further understand the nature of genes regulated by the overexpressed-*GmDR1* gene, the function of the 192 chitin-responsive, 115 buffer-responsive genes were analyzed using the GO enrichment tool in SoyBase. Upregulated and downregulated DEGs were analyzed separately. As shown in Fig. S2, classification based on cellular component GO term the up-regulated genes are overrepresented in the nucleus (90 genes) followed by either plasma membrane or extracellular region (50 genes) (Fig. S2A). The chitin-responsive upregulated genes were overrepresented in signal transduction (40 genes) and carbohydrate metabolism process (10 genes), when genes were classified based on biological processes (Fig. S2B). The downregulated DEGs were predominately classified to the cell differentiation processes. The downregulated genes were localized to the nucleus (16%), plasma membrane (14%), and cytoplasm (13%) (Fig. S2A). Based on the molecular functions of the genes, most upregulated genes (40 genes) grouped to classes that showed DNA or protein binding activities suggesting that these genes are involved in transcriptional regulation (Fig. S2C).

The 115 buffer-responsive DEGs, regulated by overexpressed-*GmDR1*, were also classified based on GO terms (Fig. S3A-C). The majority (113 genes) of the buffer-responsive DEGs were suppressed following buffer treatment in both transgenic lines (Fig. 1D). The proteins encoded by these genes were localized mostly to nuclei, plasma membrane and extracellular region (Fig. S3A) and they showed DNA binding activities based on their molecular functions (Fig. S3C).

### Identification of genes differentially regulated in response to both chitin and buffer treatments

There were only seven genes that were differentially expressed in *GmDR1*-overexpressors as compared to nontrangenic control following both chitin and buffer treatments (Fig. 1A). Three of these seven genes have shown no known functions (Table S2). The four genes with known functions include *Glyma*.*02g268200* encoding an aminocyclopropanecarboxylate oxidase involved in ethylene biosynthesis and up-regulated by pyralid feeding [45], *Glyma*.*09G131100* encoding heavy metal transport/detoxification superfamily protein responsive to cold and drought stresses [46]; *Glyma*.*16G021000* encoding a homologue of the homeobox-leucine zipper protein ATHB-12 protein induced by ABA and water stress [47], and *Glyma*.*14G171100* encoding the homologue of the transport and Golgi organization 2 protein [48].

## Discussion

The soybean GmDR1 is an integral plasma membrane protein. It is composed of 73 amino acids with two predicted transmembrane domains and one ectodomain. It is primarily a legume-specific novel protein, homologues of which are also detected in cotton, coca, and jute [22]. *GmDR1* is expressed at a low level and suppressed rapidly during attacks by pathogens and pests [21, 22]. Overexpression of *GmDR1* enhances broad-spectrum resistance against pathogens, *F. virguliforme*, SCN (*H. glycines*), and pests, spider mites (*T. urticae*, Koch), and soybean aphid (*A. glycines*, Matsumura). The chitin, a well-recognized PAMP, is present in all these four organisms, and the transcript levels of the SA- and JA-regulated defense pathway marker genes are induced in *GmDR1-*overexpressors as compared to the nontransgenic Williams 82 control line following treatment with chitin [22]. The chitin could interact directly or indirectly with GmDR1 to induce defense pathways for induction of broad-spectrum resistance. During co-evolution of soybean with its pathogens and pests, pathogens and pests were able to gain mechanisms to suppress the expression of *GmDR1* to cause susceptibility because exchange of the GmDR1 promoter with infection-inducible and strong root- and leaf-specific promoters induced enhanced pathogen and pest resistance [22].

In this study, we used chitin, because it’s a well-recognized PAMP and present in *F. virguliforme*, SCN, spider mites, and soybean aphid, against which *GmDR1-*overexpressors induced enhanced resistance. Investigation of the chitin regulated transcriptomes of the two *GmDR1*-overexpressors revealed that chitin enhances expression of defense-related genes in the two overexpressors as compared to the nontransgenic Williams 82 control (Dataset S2). To our surprise, in addition to the known genes regulating defense responses, we identified 23 genes encoding LRR domain containing proteins including ten receptor kinases and 13 NLR proteins. The transcription levels of genes encoding nine receptor kinases and all 13 NLR proteins were induced and that of one receptor kinase gene, *Glyma*.*01G062900*, was suppressed following chitin treatment among the *GmDR1*-overexpressors compared to nontransgenic control (Dataset S2).

To determine if any of the chitin-induced nine LRR receptor kinases and 13 NLR proteins may be involved in broad-spectrum immunity, we conducted RT-qPCR on RNA samples collected from roots of the *GmDR1*-overexpressors and nontransgenic Williams 82 plants infected with *F. virguliforme* and SCN. We observed that three of the nine DEGs encoding LRR receptor kinases and seven of the 13 DEGs encoding NLR receptor proteins were significantly induced in at least one of the *GmDR1*-overexpressors following *F. virguliforme* or SCN infection (Fig. 3; Table S6). The soybean Williams 82 genome carries at least 319 NLR genes [11], of which only a very few have been partially or fully characterized (*Rps11, Rps1*-k, *Rps4, Rpg1*-b, *Rsv1*) [49-53]. This work has identified several candidate NLR genes that could be involved in pathogen and pest resistance and laid a strong foundation for functional characterization of additional soybean *NLR* genes involved in immunity.

PAMP and microbe associated molecular patterns (MAMPs) are recognized by cell surface/plasma membrane-localized pattern recognition receptors (PRRs), which include receptor-like kinases (RLKs) and receptor-like proteins (RLPs). The PRR, FLAGELLIN SENSING 2 (FLS2) receptor recognizes the PAMP, bacterial flagellin flg22 and activates defense pathways in *Arabidopsis thaliana* [54]. Chitin is a PAMP widely present in many organisms. In rice, CEBiP transmembrane receptor recognizes chitin and induces defense pathways [55].

PAMPs, MAMPs, host derived damage- or danger-associated molecular patterns (DAMPs) are recognized by cell surface/plasma membrane-localized pattern recognition receptors (PRRs), which include mostly receptor kinases (RKs), receptor-like kinases (RLKs), and receptor-like proteins (RLPs). To date, 69 PRRs [56-58; Table S7] have been reported. The 29 of the 69 PRRs were identified from *Arabidopsis* (Table S7).

Twelve of the ligands recognized by these receptors are from bacteria, thirteen from fungi, two from viruses, one from aphid, and nine are host derived DAMPs. Among the 17 PRRs identified from *Solanum* species, 14 recognize bacterial ligands, one recognizes a fungal ligand, two recognize DAMPs. Among the 13 PRR from *Oryza* species, seven recognize bacterial and six fungal ligands. From *Lotus, Nicotiana*, and *Brassica* species, seven PRRs were detected that recognize bacterial ligands. On the contrary, the two PRRs from *Triticum* species are unidentified orphans that recognize fungal ligands. Only one PRR has been identified from *Glycine*, an orphan protein that recognizes a fungal beta glucan-binding protein (GBP). The 27 of the 69 identified PRRs are RLPs (39%), 21 (30%) are RKs, 16 (23%) are RLKs, and only one is LRR-RP.

Our results showed that *GmDR1* regulates several immunity-related receptor kinases and NLR receptors presumably involved in the expression of resistance against diverse pathogens and pests; and therefore, GmDR1 is most likely a key PAMP recognition receptor that induces PTI against both pathogens and pests in soybean. The GmDR1 protein, responsive to chitin and multiple pathogens and pests, is most likely a novel PRR protein and it does not fall into any of the characterized PRR classes reported earlier and represents a member of a new class of PRRs (Table S7).

Induction of a large number LRR domain containing receptor protein genes among the GmDR1-overexpressors is very intriguing. We hypothesized that only a subset of the GmDR1-induced 22 receptor protein genes including 13 *NLRs* may regulate downstream defense genes following infection against a specific pathogen or pest. The RT-qPCR data supported our hypothesis. We have shown that seven of the 13 *NLR*-type disease resistance receptor protein genes were significantly induced in at least one of the two *GmDR1*-overexpressors as compared to the nontransgenic Williams 82 control following infection with either *F. virguliforme* or SCN (Fig. 3).

In this study, we have established that the enhanced broad-spectrum disease and pest resistance among the *GmDR1* overexpressors mediated through induced expression of genes involved in plant immunity [59]. GmDR1 most likely recognizes one or more PAMPs from multiple pathogens and pests and activates multiple signaling pathways to provide soybean with immunity against pathogens and pests. It’s therefore most likely a PAMP recognition receptor for inducing PTI against several pathogen and pests; and chitin could be one of the PAMPs recognized by GmDR1. This transcriptomic study also indicates that GmDR1 may recognize other environmental ques including DAMPs as indicated by the responses of the *GmDR1* overexpressors to phosphate buffer.

Induction of nine and repression of one LRR-receptor protein kinases is also intriguing and we hypothesize that one or more of these receptor kinases may partner with GmDR1 containing only 73 amino acids during recognition of molecular patterns-specific to pathogen, pest, or tissue-damages. This study most importantly revealed that the broad-spectrum immunity induced by the overexpression of *GmDR1*could be mediated by 13 NLR proteins, a subclass of which was found to be induced by two soybean pathogens. NLRs are involved in the expression of ETI. This study indicates that NLRs could also be involved in the expression of PTI. Alternatively, GmDR1-induced NLRs may be involved in effector triggered immunity against the two pathogens. Requirement of PTI in the expression ETI has recently been reported [60]. The possible activation of ETI pathway by a pattern recognition receptor GmDR1 for PTI and/or ETI certainly very intriguing. It suggests an interdependence between PTI and ETI and priming of ETI by PTI for effective use of the cellular defense mechanisms for expression of broad-spectrum basal as well as gene- or race-specific disease resistance against multiple pathogens.

## Materials and Methods

### Soybean plant material, growing conditions and inoculation

Three soybean lines were used in this study are the non-transgenic soybean cultivar Williams 82 (W82), and two *GmDR1* transgenic lines (*GmDR1*-overexpressors). The transgenic lines DR1-136, the weak *GmDR1*-overexpressor (Promoter 2-*GmDR1*), and DR-107, the strong *GmDR1*-overexpressor (Promoter 3-*GmDR1*), were previously generated in the cultivar Williams 82 background [22]. The soybean plants were grown on standard soil in a growth chamber maintained at the constant temperature of 23°C and a photoperiod of 16 h of light/ 8 h of dark.

For the root-inoculation experiment, the seeds were sown on soil/sand mixture carrying *F. virguliforme* inocula as previously described [21, 61]. The soybean seeds of *GmDR1* overexpressors DR1-136 and DR1-107, and nontransgenic Williams 82 (W82) were sown in a 1:1 mixture of sand:soil inoculated with *F. virguliforme* inoculum at a ratio of 1 : 20:: inoculum : soil mix as reported earlier [21, 61].

For SCN inoculation, seeds were sown on soil collected from Muscatine, Muscatine County, Iowa, containing ∼50 cysts of the SCN HG type 2.5.7 (Race 5) [62]. For SCN infection, seeds were sown in cone-tainers filled with the Muscatine soil and grown in a water bath set at 27 ºC under natural light conditions for two weeks [22, 62].

### Chitin treatment

At the first trifoliate stage, two weeks following sowing, the seedlings were used for the chitin experiment. We used a modified stem-cut method for chitin (Sigma-Aldrich) treatment [63]. Half of the plants for each genotype received 0.5 mL of 100 μmol/L chitin in sodium phosphate (15 mmol/L, pH 6.5), and the second half of the plants was used as control and fed with 0.5 mL of sodium phosphate buffer (15 mmol/L, pH 6.5). All stem-cuts were fed with distilled water after chitin or phosphate buffer was depleted, and plants were maintained in the growth chamber under white light (85 μmol · m– 2 · s–1). We harvested leaf samples at 12 and 24 h post-treatment. Collected leaves were immediately placed in liquid nitrogen and stored at -80 °C for RNA isolation.

### RNA isolation and reverse transcriptase quantitative polymerase chain reaction (RT-qPCR) analysis

*F. virguliforme*-infected root samples were harvested 14 days following inoculation and used for RNA isolation. Root samples of SCN infected plants were harvested 15 days following planting and used for RNA isolation. Leaves of the soybean stem cuts 12 and 24 h following chitin treatment were used for RNA preparation. The total RNA samples were extracted from leaves or roots of nontransgenic Williams 82, DR1-136, and DR1-107 lines using the SV total RNA Isolation System that includes on-column DNase treatment (Promega, Inc., Madison, WI, USA). The quality of RNAs was assessed on agarose gel and NanoDrop microvolume spectrophotometer (Thermo Scientific, Waltham, MA, USA).

A reverse transcription kit (Superscript III First-strand synthesis SuperMix, Invitrogen by life technology) was used to synthesize first-strand cDNA from 1-2 μg total RNAs of each RNA sample. The sequences of selected 28 DEGs were retrieved from Phytozome v13 and SoyBase for designing primers [64, 65]. The primers used in the RT-qPCR are listed in Table S8. The soybean *Elongation factor 1-b* (*ELF1-b*) gene (*Glyma*.*02g44460*) was used as an internal control in RT-qPCR. The reaction was carried out in a 96-well microtiter plate on an iQ5 Biorad instrument using SYBR Green Master Mix, Applied Biosystems, by life technologies (Austin, TX). The relative expression levels of genes were evaluated using the 2−ΔΔCT method [66]. Three biological repeats, each comprising three technical replications, were conducted.

### RNA-sequencing, read mapping and transcript assembly, identification and differential expression analysis

RNA samples were sequenced at the Iowa State University DNA facility, using the HiSeq3000 and Miseq Illumina Next Generation Sequencing platforms. The analysis of RNA Seq preformed using “Tuxedo suite” that offers a set of tools for analyzing RNA-Seq data. These RNA-seq analysis tools generally fall into three categories: (i) those for read alignment; (ii) those for transcript assembly or genome annotation; and (iii) those for transcript and gene quantification [67]. We further analyzed the genes found statically significant (option “yes”) with *p* greater than the FDR after Benjamini-Hochberg correction for multiple-testing. The generated RNA-seq data set included the FPKM values for each gene. The initial analysis consisted of the comparison between Williams 82 control and each of the two transgenic lines, DR1-136, and DR1-107, 12 and 24 h following buffer and chitin treatments. To investigate the chitin induced genes and genes involved in resistance, DEGs between Williams 82 and transgenic lines were identified in each of the specific comparison detailed in Dataset S1.

Next we used Perl software (https://www.perl.org/get.html) and Microsoft excel to identify commonly up-regulated or down-regulated in both transgenic lines when compared to Williams 82 at 12 and 24 h following buffer and chitin treatments (Dataset S1). The lists of DEGs commonly upregulated and downregulated in both transgenic lines were further analyzed in the bioinformatics and evolutionary genetics program (http://bioinformatics.psb.ugent.be/webtools/Venn/), which detected the overlapping DEGs. The DEGs, up or down regulated in both transgenic lines as compared to the corresponding transcription levels in Williams 82 with an average absolute fold change ≥ 2 were retained for subsequent analysis. The final number of DEGs was therefore reduced to 115 from 143 for DEGs in buffer treatment and to 192 from 289 for DEGs following chitin treatment (Datasets S2 and S3).

### MapMan and Gene ontology (GO) annotation analyses

To determine gene function, we used the plant-specific visualization tool, MapMan, to identify the DEGs that are involved in specific pathways. The MapMan software was obtained from https://mapman.gabipd.org/ to visualize the genes by using the transcriptomic data of this study. The mapping files from the MapMan Store (http://mapman.gabipd.org/web/guest/mapmanstore) were used. The genome release version 3.6 was used to transfer soybean genome sequence to MapMan ontology (file name: Map files/Gmax_189.txt), and then made the gene name transferred into gene name of Gmax_275 version according to the file (Gmax_275_Wm82.a2.v1.synonym.txt) downloaded from Phytozome (https://phytozome.jgi.doe.gov/pz/portal.html). The automated annotations using the soybean sequences allowed the assignment of more than 54,505 gene classifications into 35 BINs. Amino acid sequences of proteins encoded by the DEGs were extracted from Phytozome [64] and SoyBase [65]. To identify the conserved domains and annotation descriptions, the amino acid sequences were blasted in NCBI. Gene ontology (GO) annotation analyses of DEGs were conducted based on their molecular functions, biological processes and cellular components using SoyBase GO term enrichment tool.

## Acknowledgments

This project was supported by USDA-NIFA (grant no. 2013-68004-20374) and Iowa Soybean Association.

